# Dataset of emerging contaminants in surface water, bottom water, porewater, and sediment: Urban and aquaculture impacts in the central and southern coast of Chile

**DOI:** 10.1101/2024.04.30.591869

**Authors:** Pedro A. Inostroza, Yolanda Soriano, Eric Carmona, Martin Krauss, Werner Brack, Thomas Backhaus, Renato A. Quiñones

**Affiliations:** Institute for Environmental Research, RWTH Aachen University, Aachen, Germany; Department of Biological and Environmental Sciences, University of Gothenburg, Gothenburg, Sweden; Food and Environmental Safety Research Group of the University of Valencia (SAMA-UV), Desertification Research Centre (CIDE) CSIC-GV-UV, Valencia, Spain; Department Exposure Science, Helmholtz Centre for Environmental Research – UFZ, Leipzig, Germany; Department of Evolutionary Ecology and Environmental Toxicology, Goethe University Frankfurt/Main, Frankfurt/Main, Germany; Interdisciplinary Center for Aquaculture Research (INCAR), University of Concepcion, Concepción, Chile

**Keywords:** Target screening, pesticides, pharmaceuticals, marine pollution, LVSPE, marine pollution

## Abstract

Synthetic organic chemicals, including pesticides, pharmaceuticals, and industrial compounds, pose a growing threat to marine ecosystems as they enter through a variety of pathways, including direct discharges of wastewater (untreated or treated) from industrial, agricultural, and urban sources. Additionally, runoff from residential and agricultural land, as well as inland waterways, transport these chemicals to coastal zones. Despite their potential impact, data on the co-occurrence of these contaminants in the marine environment remains limited. Such information is critical for assessing coastal chemical status, establishing environmental quality benchmarks, and conducting comprehensive environmental risk assessments. In this study, we describe a multifaceted monitoring campaign targeting pesticides, pharmaceuticals, and industrial chemicals along the central-south coast and in northern Patagonia, Chile. Surface water, bottom water, porewater, and adjacent sediment samples were collected for analysis. Our results show the detection of up to 83 chemicals in surface water, 71 in bottom water, 101 in porewater, and 244 in sediments. To enhance data utility, we provide valuable information on the mode of action and molecular targets of the identified chemicals. This comprehensive dataset contributes to defining pollution fingerprints in coastal areas of the Global South, including remote regions in Patagonia. It serves as a critical resource for future research, policymaking, and the advancement of environmental protection in these regions.

## SPECIFICATIONS TABLE

42 SPECIFICATIONS TABLE
|  |  |
| --- | --- |
| Subject | Pollution |
| Specific subject area | Synthetic organic pollutants occurring in coastal areas |
| Type of data | Table<br>Analyzed, Filtered |
| Data collection | The data were acquired via gas and liquid chromatography-high-resolution mass spectrometry. Target screening was conducted for 861 chemicals using an UltiMate 3000 LC system (Thermo Scientific, Germany) coupled to a hybrid quadrupole-Orbitrap MS (Q Exactive Plus, Thermo Scientific, Germany) with a heated electrospray ionization (ESI) source. A retrospective analysis was applied to 150 of the 861 target chemicals. More hydrophobic analytes, comprised of 36 chemicals, were re-evaluated using a TRACE 1310 GC system (Thermo Scientific, Germany) coupled to a quadrupole-Orbitrap MS (Q Exactive, Thermo Scientific, Germany) equipped with a Thermal Desorption Unit (TDU-2; Gerstel, Mülheim, Germany) and a cooled injection system (CIS; Gerstel). |
| Data source location | Data were stored: Institute for Environmental Research, RWTH Aachen University<br>City/Town/Region: Aachen, Nordrhein-Westfalen<br>Country: Germany<br>Data were collected: Latitude and longitude (WGS84):<br>Coliumo (36.531639 S 72.951917 W),<br>Tubildad (42.124361 S 73.470944 W),<br>Aucar (42.162694 S 73.480833 W)<br>Montolin (42.207639 S 73.390611 W) |
| Data accessibility | Repository name: zenodo<br>Data identification number: 10.5281/zenodo.11080740<br>Direct URL to data: https://zenodo.org/records/11080740 |

## VALUE OF THE DATA

- We report environmental concentrations of emerging chemicals, including pesticides, pharmaceuticals, and industrial chemicals, in coastal areas of central-southern Chile and Northern Patagonia, Chile.
- The dataset includes multi-compartment measurements (surface water, bottom water, porewater, and sediments) from each sampling site.
- The reported data can be used for the definition of pollution profiles and for the establishment of environmental quality standards.
- The data can be used by environmental risk assessors for risk prediction and by local authorities to develop mitigation strategies and environmental surveillance programs.

## DATA DESCRIPTION

The dataset in this study originates from surface water, bottom water, porewater, and superficial sediment samples collected from four distinct sites affected by anthropogenic activities in the central-southern coast and northern Patagonia of Chile.

The dataset is reported in tabular format and is available in both Rdata (RDS) and tab-separated values (TSV) formats. The dataset can be accessed at Inostroza et al. [1]. For each reported substance, the dataset includes essential identifiers such as the CAS Registry Number (CAS RN), the International Chemical Identifier (InChI), and its hashed InChIKey. To complement the dataset, chemical classes and sub-classes, based on their intended use, are included (e.g., Class 1: pharmaceutical, Class 2: antihypertensive, Class 3: beta blocker). Additionally, the mode of action (MoA) information (broad and specific) as well as molecular target sites of each chemical was retrieved from Kramer et al. [2]. The RDS and TSV files contain the columns defined in Table 1, while Table 2 provides an overview of the micropollutants detected and quantified.

**Table 1.** Columns and its respective description in the dataset.

| Columns |  |
| --- | --- |
| Chemical_name | Name of the chemical |
| CAS_number | CAS Registry Number use as chemical identifier |
| InChIKey | Textual identifier for chemical substances |
| Screening_method | Screening platform used for analysis |
| Class_1 | Use category (e.g., pharmaceutical, pesticide, biocide, etc.) |
| Class_2 | Sub-use category (e.g., antibiotic, herbicide, plasticizer, etc.) |
| Class_3 | Sub-use category (e.g., benzodiazepine, organophosphate, etc.) |
| MoA_broad | Mode of action of the chemical |
| MoA_specific | Specific Mode of action of the chemical |
| Molecular_target | Molecular target known to interact with the chemical |
| Matrix | Environmental compartment |
| Sites | Sampling sites |
| MDL | Method detection limit in ng/L |
| Quant | Measured environmental concentrations in ng/L |
| units | Unit of the measurements (ng/L for water and ng/g for sediment) |

**Table 2.**
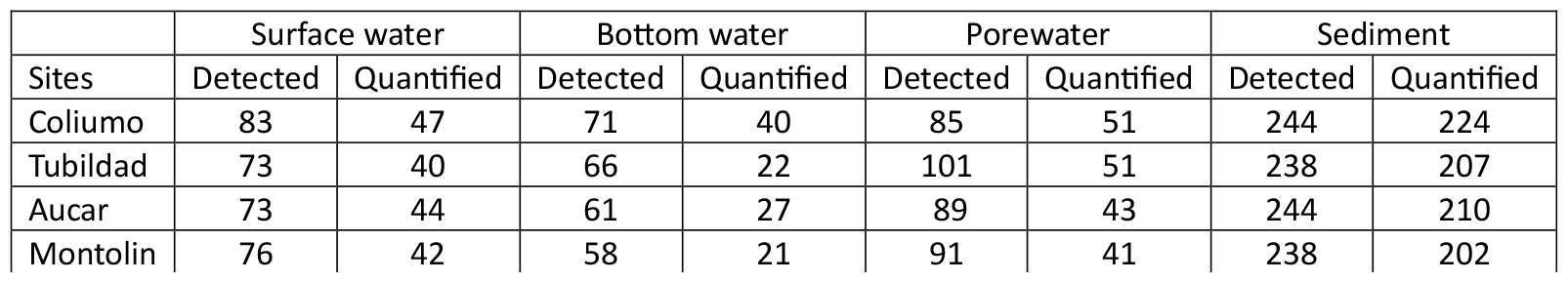
Summary of detected and quantified chemicals per environmental matrix.

## EXPERIMENTAL DESIGN, MATERIALS AND METHODS

### Sampling design and sample collection

Samples were collected from four sampling sites along central-southern coast of Chile and northern Patagonia’s coast, Chile, in October 2021 (spring season) (Figure 1A). Sampling sites were selected based on the type of land use in the surrounding area. This allowed us to characterise coastal sites with different levels of anthropogenic impact.

**Figure 1.**
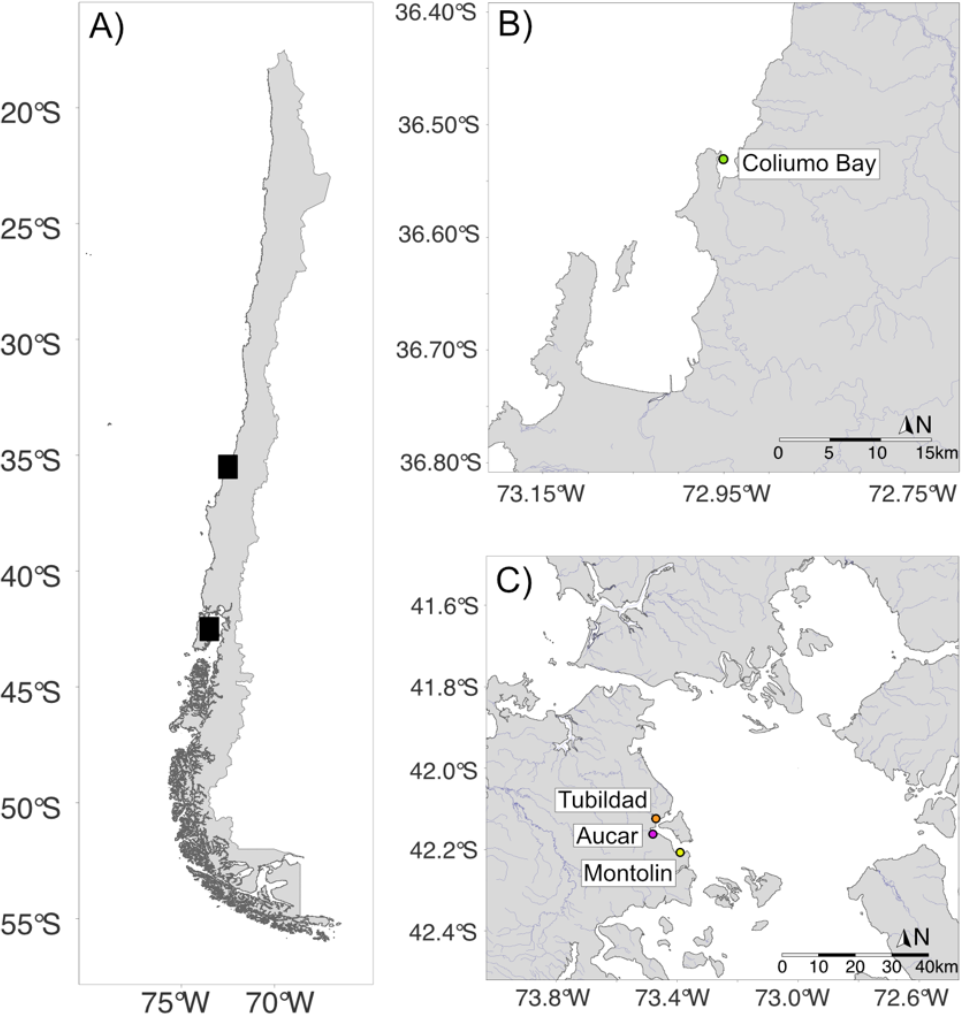
A) Location of sampling sites along the Chilean coast. B) Sampling site in Coliumo Bay. C) Sampling sites in the northern Patagonia (Caucahue Channel). All sampling sites are color-coded.

Coliumo Bay, a small bay in south-central Chile, is home to approximately 7,500 people and serves as a popular summer tourist destination. Additionally, the bay supports a small fishing fleet and receives both treated and untreated wastewater discharges. It is important to note that there is no salmon farming in Coliumo Bay or the surrounding area (Figure 1B).

Tubildad, Aucar, and Montolin are located in northern Patagonia, specifically within the Caucahue Channel (Figure 1C). This channel is subject to significant influence from aquaculture activities, including 37 mussel farms, 13 salmon farms, and five seaweed farms [3]. It is also impacted byagricultural activities, along with the discharge of untreated and treated urban and industrial wastewater. Further details on the sampling locations and geographical coordinates of the WWTPs are given in Table 3.

**Table 3.** Additional sampling site information. Geographic coordinates in decimal degree (WGS84).

| Sites | Latitude (S) | Longitude (W) | WWTP | Type of treatment |
| --- | --- | --- | --- | --- |
| Coliumo | 36.531632 | 72.951917 | Dichato | Activated sludge |
| Tubildad | 42.124361 | 72.470944 | - |  |
| Aucar | 42.162694 | 73.480833 | Quemchi | Activated sludge |
| Montolin | 42.207639 | 73.390611 | - |  |

Surface water samples were collected from 2 meters below the surface using a Niskin bottle (20 L, General Oceanics). Water was transferred to 20 L glass bottles, previously cleaned with methanol, and filtered using an on-site large-volume solid phase extraction (LVSPE) device (MAXX Mess-und Probenahmetechnik GmbH, Rangendingen, Germany). The LVSPE enabled the filtering of 50 L per sampling and has been previously employed in marine sampling [4]. A detailed description of the LVSPE sampler, method development, and extraction recoveries can be found in Schulze et al. [5]. Cartridge preparation, conditioning, and extraction are also explained in detail in Schulze et al. [5]. Subsequent liquid chromatography (LC) and gas chromatography (GC) analyses followed established analytical protocols [4,6,7]. To ensure sample integrity, the remaining extracts were stored at −20 °C for future bioassays.

Surface sediment samples were collected using a cylindrical piston corer equipped with a 6 cm diameter plastic core tube (USC 06000, UWITEC GmbH, Austria). The top 4-5 cm were sliced using a stainless-steel cutting plate previously cleaned with methanol. Samples were immediately stored in aluminum boxes in the dark at -20 °C in a portable freezer until extraction.

Bottom water and porewater were collected using the cylindrical piston corer. The corer was predrilled at 3 cm intervals and taped before each sampling. Rhizons (0.45 µm pore size, Rhizosphere Research Products, Netherlands) were inserted at 3 cm intervals in the predrilled holes to collect bottom and porewater samples. These samples were stored in 5 mL brown vials with headspace (i.e., leaving some air in the vial) and kept at -20°C.

### Bottom, porewater, and sediment preparation and extraction procedures

First, bottom and porewater samples were filtered using Whatman GF/F 55 mm filters (GE Healthcare). Solid-Phase Extraction (SPE) was performed on these samples using Chromabond HR-X cartridges (6 mL, 200 mg sorbent, Macherey-Nagel, Düren, Germany) with a Promochrom SPE-03 automated device. SPE process blanks were included in parallel.

Before sample loading, SPE cartridges were conditioned with 5 mL of ethyl acetate (10 mL/min) followed by 5 mL of methanol (10 mL/min). After sample extraction, cartridges were washed with 1 mL of ultra-pure water and dried for approximately 90 minutes using N2. Sample extracts were eluted with the following solvent fractions at a flow rate of 5 mL/min: 5 mL ethyl acetate, 5 mL methanol, 4 mL methanol with 1% formic acid, and 4 mL methanol with 2% 7N ammonia. The extracts were evaporated under nitrogen flow to approximately 500 µl, filtered (PTFE, 0.2 µm pores size, 13 mm diameter), and then evaporated to dryness. Extract residues were reconstituted in 0.5 mL of methanol.

Sediment samples were extracted by Pressurized Liquid Extraction (PLE) with a mixture of acetone and ethyl acetate (1:1 v/v) using an Accelerated Solvent Extraction (ASE) 200 device (Dionex). The raw extracts were concentrated under a nitrogen stream and the solvent was exchanged for dichloromethane. The extracts were then evaporated to near dryness using a XcelVap nitrogen evaporator, and finally adjusted to 0.5 mL with dichloromethane. The extracts were transferred to 2 mL glass vials and stored at -20°C until purification.

Sediment cleanup was performed by flash chromatography using an Agilent 1200 binary pump and a pre-packed chromatography column (Chromabond Flash RS 4 g SiOH column, 10.6 cm × 12.4 mm, Macherey-Nagel, Düren, Germany). The column was first conditioned with dichloromethane and the extracts were eluted with dichloromethane and methanol, which were collected in separate fractions.

For the GC-HRMS analysis, half of the dichloromethane fraction was concentrated under a nitrogen stream and re-dissolved in ethyl acetate. The second half of the dichloromethane fraction was combined with half of the methanol fraction, concentrated under a nitrogen stream and re-dissolved in methanol for LC-HRMS analysis. Processing blanks were prepared with hydromatrix and processed the same way as the samples. All samples were filtered (pore size 45 μm) and stored in a freezer at - 20 °C until analysis.

### Target chemical screening in LC-HRMS and GC-HRMS

Monitoring of emerging chemicals, such as pesticides, pharmaceuticals, and industrial chemicals, is limited in coastal areas of South America, particularly in Chile [8,9]. Therefore, we employed a target list of 861 chemicals, based primarily on those commonly detected in European freshwater systems. However, it’s important to note that the same target list was previously used for surface waters collected along the Swedish coast [4]. Furthermore, the classification system for these chemical classes is predominantly European/German-centric.

The target screening was conducted using an UltiMate 3000 LC system (Thermo Scientific) coupled with a hybrid quadrupole-Orbitrap MS (QExactive Plus, Thermo Scientific) featuring a heated electrospray ionization (ESI) source. A retrospective analysis, as outlined by Muschket et al. [10], was applied to 150 out of the 861 target chemicals. Furthermore, an additional evaluation using a TRACE 1310 GC system (Thermo Scientific) coupled with a hybrid quadrupole-Orbitrap MS (QExactive GC, Thermo Scientific) was performed for the more hydrophobic analytes (36 chemicals). This GC system was equipped with a Thermal Desorption Unit (TDU-2; Gerstel, Mülheim, Germany) and a cooled injection system (CIS; Gerstel). More detailed information on the settings for LC-HRMS and GC-HRMS can be found in [4,7].

ProteoWizard (V 2.1.0) was used to convert LC-HRMS raw data into mzML format [11]. Subsequently, peak detection, sample alignment, and target compound annotation were performed using MZmine (V 2.40.1) [12] as detailed in Beckers et al. [13]. We used the R package {MZquant} (V 0.7.22) to perform blank correction, calibration, and then quantification of the annotated target compounds. Blank peak elimination and blank intensity thresholds were calculated according to Machate et al. [7].

For the quantification of GC-HRMS detected compounds, the software TraceFinder 4.1 (Thermo Scientific) was used for further evaluation. Method-matched calibration standards were used in a series ranging from 0.5 to 5000 ng/L. These calibration standards were treated in the same way as the samples. The target compounds were quantified using the internal standards with the nearest retention time following Nanusha et al. [6] and Machate et al. [7]. The method detection limits (MDLs) were determined according to US-EPA procedure [14].

## LIMITATIONS

The chemical target list for micropollutant analysis in this study was primarily based on chemicals commonly detected in streams and rivers in European contexts. Besides, the classification of chemical classes is predominantly European/German-centric. The classification of modes and mechanisms of action relates for the vast majority of chemicals to the intended target organisms or close phylogenetic relatives.

## ETHICS STATEMENT

The proposed data does not involve any human subjects, animal experiments, or data collected from social media platforms.

## CRediT AUTHOR STATEMENT

### Pedro A. Inostroza

Conceptualization, Methodology, Formal analysis, Investigation, Data Curation, Writing - Original Draft, Writing - Review & Editing, Visualization. **Yolanda Soriano**: Methodology, Validation, Formal analysis, Writing - Review & Editing. **Eric Carmona**: Methodology, Validation, Formal analysis, Writing - Review & Editing. **Martin Krauss**: Methodology, Validation, Investigation, Writing – Review. **Werner Brack**: Investigation, Resources, Writing - Review & Editing. **Thomas Backhaus**: Resources, Writing - Review & Editing, Funding acquisition. **Renato A. Quiñones**: Conceptualization, Resources, Investigation, Writing - Review & Editing. All authors have read and agreed to the published version of the manuscript.

## ACKNOWLEDGEMENTS

We thank Oliver Alarcon and Vicente Villalobos for fieldwork support, Margit Petre and Jörg Ahleim (UFZ) for LVSPE cartridge preparations. This work was supported by the FRAM Centre for Future Chemical Risk Assessment and Management at the University of Gothenburg and the Interdisciplinary Center for Aquaculture Research (INCAR, FONDAP-ANID, Grants 1522A0004). The QExactive Plus LC-HRMS and GC-HRMS used at UFZ are part of the major infrastructure initiative CITEPro (Chemicals in the Terrestrial Environment Profiler) funded by the Helmholtz Association.

